# Connecting the dots: Harnessing dual-site transcranial magnetic stimulation to assess the causal influence of medial frontal areas on the motor cortex

**DOI:** 10.1101/2022.01.18.476729

**Authors:** Cécilia Neige, Pierre Vassiliadis, Abdelkrim Ali Zazou, Laurence Dricot, Florent Lebon, Thomas Brees, Gerard Derosiere

**Affiliations:** Université Bourgogne Franche-Comté, INSERM UMR1093-CAPS, UFR des Sciences du Sport, F-21078, Dijon, France; Université Claude Bernard Lyon 1, CNRS, INSERM, Centre de Recherche en Neurosciences de Lyon CRNL U1028 UMR5292, PsyR2 Team, F-69500, Bron, France; Centre Hospitalier le Vinatier, F-69500 Bron, France; Institute of Neuroscience, Université Catholique de Louvain, 1200, Brussels, Belgium; Defitech Chair for Clinical Neuroengineering, Neuro-X Institute (INX) and Brain Mind Institute (BMI), École Polytechnique Fédérale de Lausanne (EPFL), 1202, Geneva, Switzerland; Université Claude Bernard Lyon 1, CNRS, INSERM, Centre de Recherche en Neurosciences de Lyon CRNL U1028 UMR5292, Impact Team, F-69500, Bron, France

**Keywords:** Corticospinal excitability, effective connectivity, medial orbitofrontal cortex, supplementary motor area, magnetic functional imagery.

## Abstract

Dual-site transcranial magnetic stimulation (TMS) has been widely employed to investigate the influence of cortical structures on the primary motor cortex (M1). Here, we leveraged this technique to probe the causal influence of two key areas of the medial frontal cortex, namely the supplementary motor area (SMA) and the medial orbitofrontal cortex (mOFC), on M1. We show that SMA stimulation facilitates M1 activity across short (6 and 8 ms) and long (12 ms) inter-stimulation intervals, putatively recruiting cortico-cortical and cortico-subcortico-cortical circuits, respectively. Crucially, magnetic resonance imaging revealed that this facilitatory effect depended on a key morphometric feature of SMA: individuals with larger SMA volumes exhibited more facilitation from SMA to M1. Notably, we also provide evidence that the facilitatory effect of SMA stimulation at short intervals did not arise from spinal interactions of volleys descending simultaneously from SMA and M1. On the other hand, mOFC stimulation moderately suppressed M1 activity at both short and long intervals, irrespective of mOFC volume. These results suggest that dual-site TMS is an interesting tool to study the differential influence of SMA and mOFC on M1 activity, paving the way for the multi-modal assessment of these fronto-motor circuits in health and disease.

**Key points:**

- Dual-site TMS has been widely employed to investigate effective connectivity between cortical structures and the primary motor cortex (M1).
- Here, we probed the causal influence of the supplementary motor area (SMA) and the medial orbitofrontal cortex (mOFC) on M1 activity.
- SMA stimulation facilitates M1 activity at both short and long inter-stimulation intervals; this facilitatory effect is related to SMA volume.
- mOFC stimulation moderately suppresses M1 activity, independent of mOFC volume.
- The findings pave the way for multi-modal assessment of fronto-motor circuits in health and disease.

## Introduction

The execution of most volitional actions relies on pyramidal cells located in the primary motor cortex (M1), that project down to the spine and connect with peripheral motoneurons. This so-called corticospinal pathway is under the constant influence of distributed areas of the cerebral cortex, that rely on effective connectivity to either facilitate or suppress M1 activity and, ultimately, exert control over behavior. As such, two key areas of the medial frontal cortex– the supplementary motor area (SMA) and the medial orbitofrontal cortex (mOFC) – are known to play a central role in human behavior, being involved in processes as diverse as motor planning and learning (Carlsen et al., 2015; Makoshi et al., 2011; Neige et al., 2018; Vassiliadis et al., 2023), decision-making (Fellows, 2007; Klein-Flügge et al., 2016; Derosiere et al., 2018; Loh & Rosenkranz 2022) and inhibitory control (Aron et al., 2007; Boy et al., 2010; Hu & Li 2012). SMA and mOFC project to M1 via both cortico-cortical and cortico-subcortico-cortical circuits, providing candidate pathways through which they might implement these processes. Crucially, however, tools to probe their effective influence on M1 remain scarce at present.

In humans, a specific type of transcranial magnetic stimulation (TMS) protocol – called dual-site paired-pulse TMS (ppTMS) – allows probing the effective connectivity between specific cortical areas and M1 (for recent reviews, see Derosiere & Duque 2020; Neige et al., 2021). As such, the excitability of the corticospinal pathway can be quantified by recording of motor-evoked potentials (MEPs), which can be elicited in muscles by applying single-pulse TMS over the contralateral M1. The amplitude of MEPs provides a global readout of corticospinal excitability, reflecting the simultaneous influence of multiple brain structures projecting to M1 (Bestmann & Duque 2016; Di Lazzaro et al., 2018). Dual-site ppTMS allows to isolate the influence of a targeted cortical area on M1. In such protocols, a first, conditioning stimulation is used to pre-activate the targeted area, while a second, test stimulation is applied over M1 with another coil to elicit a MEP and assess the nature of the influence (*i.e.*, facilitatory or suppressive) of the pre-activated area on corticospinal excitability. Potentiation of conditioned MEP amplitudes (*i.e.*, relative to unconditioned MEPs) reflects a facilitatory influence of the pre-activated area, whereas a reduction of conditioned MEPs reflects a suppressive effect. Interestingly, varying the inter-stimulation interval allows to probe different circuits, with short inter-stimulation intervals (*e.g.*, between 4 and 8 ms) recruiting cortico-cortical circuits preferentially, and longer ones (*e.g.*, higher than 10 ms) recruiting more indirect circuits presumably funneling through subcortical structures (Neubert et al., 2010).

Over the past two decades, dual-site ppTMS has been widely used in humans, with studies probing the causal influence of several areas of the premotor cortex (Koch et al., 2006; Davare et al., 2008), of the lateral prefrontal cortex (Neubert et al., 2010; Wang et al., 2020) and of the parietal cortex on M1 (Koch & Rothwell 2009; Koch et al., 2009; Lebon et al., 2012; Vesia et al., 2017; Allart et al., 2019). However, several issues remain open, in particular regarding the use of this approach on some areas of the medial frontal cortex – including SMA and mOFC.

To date, the few ppTMS studies targeting SMA have mostly focused on short inter-stimulation intervals (6 to 8 ms, Arai et al., 2011, 2012; Green et al., 2018; Rurak et al., 2021). Given this, there is currently a lack of data on the nature of the influence (*i.e.*, facilitatory *vs*. suppressive) of SMA stimulation on M1 activity when probed with longer inter-stimulation intervals (*i.e.*, 10 to 15 ms). In fact, the SMA projects to M1 through multiple cortico-subcortico-cortical circuits (Nachev et al., 2008), some of which exert a net facilitatory influence on motor activity (*e.g.*, the direct basal ganglia pathway) and some of which play a suppressive role (*e.g.*, the indirect and hyperdirect pathways). Interestingly, ppTMS studies focusing on the preSMA – *i.e.*, another key area of the medial frontal cortex – with intervals of 12 ms, reported a potentiation of conditioned MEP amplitudes, which strongly covaried with white matter density in preSMA-basal ganglia-M1 circuits (Mars et al., 2009; Neubert et al., 2010). Taken together, these two sets of findings suggest that, when applied over preSMA at such intervals, ppTMS recruits circuits that have a facilitatory influence on M1 and funnel through the basal ganglia. A first goal of the present study is to shed light on to the influence of SMA stimulation on M1 at such intervals, by testing the idea that SMA-originating circuits bear a similar facilitatory influence on motor activity.

As such, ppTMS studies targeting SMA with short inter-stimulation intervals (6 to 8 ms) have reported a potentiation of MEP amplitudes, which has been assumed to reflect the operation of cortico-cortical, facilitatory circuits from SMA to M1 (Arai et al., 2011, 2012; Green et al., 2018; Rurak et al., 2021). Indeed, this assumption seems consistent with animal studies showing that SMA has direct glutamatergic projections to M1 (Muakkassa & Strick 1979; Luppino et al., 1993) and that electrical stimulation of SMA neurons elicits responses in M1 with short latencies, of about 4 ms (Aizawa & Tanji 1994; Tokuno & Nambu 2000).

Importantly though, other observations challenge the validity of this assumption. Indeed, like M1, SMA has pyramidal cells that project to the spine (Dum & Strick 1996) and, in certain contexts, unique stimulation of SMA with single-pulse TMS can evoke MEPs (Spieser et al., 2013; Entakli et al., 2014), suggesting that these pyramidal cells can also recruit motoneurons. Thus, it is possible that the MEP potentiation reported in ppTMS studies using short intervals reflects the summation of volleys descending from SMA and M1 and converging at close times on motoneurons. In other words, it is currently unclear whether this potentiation can be taken as a pure measure of effective connectivity between SMA and M1 or not. Addressing this issue is fundamental for any investigation targeting motor areas (*e.g.*, the dorsal or the ventral premotor cortex; Davare et al., 2009, Koch et al., 2006), which, for the most part, present corticospinal projections (Dum & Strick, 1991). This is the second goal of the current study. Specifically, we tested the effect of SMA conditioning on MEP amplitudes using a very short inter-stimulation intervals of 1 ms. The rationale here is that a 1 ms interval would be too short for a MEP potentiation to result from the recruitment of cortico-cortical circuits (Aizawa & Tanji, 1994; Tokuno & Nambu, 2000); any potentiation occurring at this interval would instead provide evidence for a summation of neural inputs occurring at the spinal level.

As mentioned above, mOFC is another major area of the medial frontal cortex (Amodio & Frith, 2006). Strikingly though, ppTMS has never been used to probe the influence of this area on M1, potentially because of the presumed difficulty of reaching it with magnetic fields. Hence, it is currently unknown whether TMS could be exploited to probe effective connectivity between mOFC and M1 and, if so, what would be the nature of the influence of the recruited circuits on motor activity. In fact, while former investigations on caudal areas of the medial frontal cortex (*i.e.*, SMA and preSMA) generally reported a facilitatory influence on M1, ppTMS studies on more rostral areas of the frontal lobe, such as the dorsolateral PFC, revealed the operation of suppressive circuits (Wang et al., 2020). A third goal of the present study is to test the feasibility of using ppTMS to probe effective connectivity between mOFC and M1 and, in turn, to determine the influence of mOFC stimulation on M1 activity.

Finally, previous studies utilizing dual-site TMS have consistently observed variations in effective connectivity between cortical areas and M1 (Ferbert et al., 1992; Brown et al., 2019; Wang et al., 2020; Rurak et al., 2021). However, the underlying neurophysiological mechanisms responsible for this inter-individual variability have remained elusive. In fact, single-pulse TMS studies have demonstrated how individual morphometric features of M1, such as cortical thickness, gray matter volume, and microstructural properties of cerebral white matter, can significantly influence cortical excitability (Klöppel et al., 2008; List et al., 2013). Therefore, in the current study, we exploited structural MRI to characterize the relationship between the morphometric features of SMA and mOFC, particularly their cortical volume, and the facilitatory/suppressive effect of their stimulation on M1 activity.

Overall, the current study addresses four main goals. The first one is to provide insight into the influence of SMA stimulation on M1 activity with long inter-stimulation intervals, which are thought to recruit cortico-subcortico-cortical circuits. As a second objective, we aim to clarify whether the MEP potentiation reported in studies targeting SMA and M1 with short intervals can be taken as a pure measure of cortico-cortical connectivity between these areas or if it could in part reflect the summation of volleys descending from those on motoneurons. Moreover, we seek to test the feasibility of exploiting ppTMS to probe effective connectivity between mOFC and M1 and, relatedly, to determine the influence of mOFC stimulation on motor activity. As a last goal, we also investigated whether these TMS-based measures of fronto-motor connectivity were associated to the individual volume of SMA and mOFC.

## Methods

### Ethical approval

The protocol was approved by the institutional review board of the Catholic University of Louvain (number 2018/22MAI/219) and complied with the latest version of the Declaration of Helsinki, except for registration in a database.

### Participants

Twenty healthy subjects participated in the current study (11 females; mean age 26.9 years ± 5.4; right-handed as assessed by the Edinburgh Handedness Inventory (Oldfield, 1971)). Subjects were recruited from the Research Participant Pool at the Institute of Neuroscience of the Catholic University of Louvain (Brussels, Belgium). None of them had any neurological disorder, history of psychiatric illness or drug or alcohol abuse, or presented any contraindication to TMS (Rossi et al., 2009, 2021).

### MRI data acquisition

Anatomic sequence was acquired at the Cliniques Universitaires Saint-Luc (UCLouvain, Belgium) using a 3T head scanner (Signa™ Premier, General Electric Company, USA) equipped with a 48-channel coil.) A three-dimensional (3D) T1-weighted data set encompassing the whole brain was selected to provide detailed anatomy (1 mm³) thanks to a MPRAGE sequence (inversion time = 900 msec, repetition time (TR) = 2238.92 msec, echo time (TE) = 2.96 msec, flip angle (FA) = 8°, field of view (FOV) = 256*256 mm², matrix size = 256*256, 170 slices, slice thickness = 1 mm, no gap). Patients, instructed to remain still, were positioned comfortably in the coil and fitted with soft earplugs.

### Transcranial magnetic stimulation protocol

#### Coil locations

Dual-site ppTMS involves applying a test stimulation (TS) with one coil over M1 preceded, in a certain proportion of trials, by a conditioning stimulation (CS) delivered with another coil over an area of interest. Our aim here was to investigate intra-hemispheric influences of SMA and mOFC on M1. To do so, we applied both stimulations over the left, dominant hemisphere, with the TS administered over the left M1 and the CS targeting either the left SMA or the left mOFC in separate blocks of trials. TS and CS were delivered with two small figure-of-eight coils (Magstim D25-Alpha model; wing internal diameter: 35 mm) connected to two monophasic Magstim stimulators (200^2^ and Bistim^2^ stimulators; Magstim, Whitland, Dyfed, UK).

#### M1 coil

The M1 coil was placed tangentially to the scalp with the handle pointing backward and laterally at a 45° angle away from the midsagittal line, resulting in a postero-anterior current flow within the cortex (Rossini et al., 1994; Rossi et al., 2009; Derosiere et al., 2015, 2020). To define the optimal site for M1 stimulation (*i.e.*, the so-called “hotspot”), we relied on markers disposed on an electroencephalography (EEG) cap fitted on the participant’s head (Zrenner et al., 2018; Derosiere et al., 2019). Of note, due to the anatomical proximity of left M1 and left SMA, in a number of subjects (n = 12 / 20), the position of the M1 coil had to be slightly adjusted when the SMA coil was settled over the scalp; this position was exploited for TS during SMA blocks. Hence, we defined two different positions for M1 stimulation: the real hotspot and an adjusted hotspot.

To find the real hotspot, we first applied the stimulation with the center of the M1 coil over the C3 location of the EEG cap (*i.e.*, corresponding to the left M1 area; Derosiere et al., 2018; Alamia et al., 2019), in the absence of the second coil on the head. Stimulation intensity was increased until consistent MEP responses were obtained in the right first dorsal interosseous (FDI) muscle at this location. We then moved the coil by steps of ∼ 0.5 cm around this location in both the rostrocaudal and the mediolateral axes. Stimulation was applied with the previously defined intensity at each new location, and MEP amplitudes were visually screened. The real hotspot was defined as the location at which the largest and most consistent MEP amplitudes could be obtained (Rossini et al., 1994; Neige et al., 2018; Derosiere et al., 2019). The coil was then held at this location, and the edges of its shape were marked on tapes disposed on the EEG cap (Vassiliadis et al., 2020; Geers et al., 2021; Neige et al., 2022; Wilhelm et al., 2022). These marks allowed us to localize the real hotspot at any required time during the session. To determine the adjusted hotspot, we first positioned the SMA coil over the head and then reproduced the same procedure as described above, while trying to fit the two coils over the head. Once the largest MEP amplitudes obtained, the two coils were held at their respective locations, and the edges of the M1 coil were marked on the EEG cap. These marks allowed us to localize the adjusted hotspot during SMA blocks.

#### SMA and mOFC coils

To ensure that the coil exploited for the CS was precisely targeting SMA and mOFC in each subject, the T1-weighted images were used for neuronavigation. Precisely, we determined the SMA and mOFC locations on the individual images using MNI coordinates in a dedicated software (Visor 2.0 Advanced NeuroTechnologies, Enschede, Netherlands). The locations were finally used during the experiment, in which we relied on head and coil trackers as well as a 3D tracking device to coregister the position of the SMA/mOFC coil with the individual MRI.

The MNI coordinates exploited to Initially localize SMA and mOFC were x = -8, y = -9, z = 77 and x = − 7, y = 71, z = -4, respectively (Codol et al., 2020). These two locations were then slightly adjusted for each subject using the Visor software, so that they corresponded to the point where the scalp-to-cortex distance was minimal. Following this procedure, the MNI coordinates for SMA and mOFC locations were x = -7.9 ± 0.3, y = -7.4 ± 0.8, z = 82.9 ± 1 and x = -9.4 ± 0.7, y = 72.6 ± 0.5, z = 8.2 ± 1.7, respectively (mean ± standard error (SE) of the group; see Figure 1 and Table 1 for group-averaged and individual MNI coordinates, respectively).

**Figure 1:**
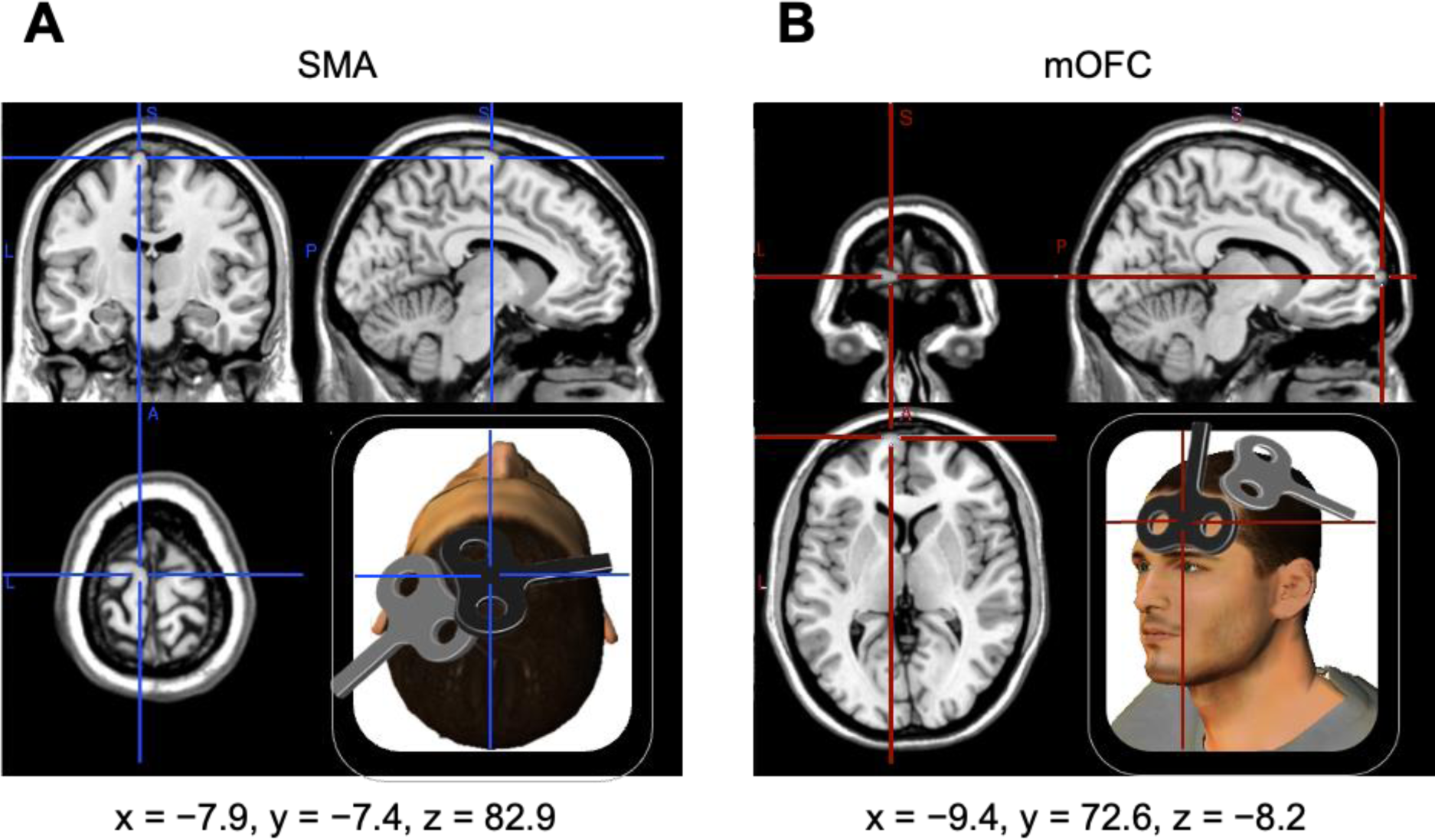
Localization of the sites of stimulation and illustration of the coil positioning for SMA (A) and mOFC (B) targets. The group-averaged MNI coordinates are illustrated on a standard MRI template (coronal, sagittal and axial views) using MRIcron software (v1.0.20190902 http://www.mricro.com).

**Table 1:** Individual MNI coordinates for SMA and mOFC target locations. The MNI coordinates exploited to initially localize SMA and mOFC were x = -8, y = -9, z = 77 and x = − 7, y = 71, z = -4, respectively. These two locations were then slightly adjusted for each subject using the Visor software, so that they corresponded to the point where the scalp-to-cortex distance was minimal.

| Subject | SMA coordinates |  |  | mOFC coordinates |  |  |
| --- | --- | --- | --- | --- | --- | --- |
|  | x | y | z | x | y | z |
| **1** | -6 | -7.7 | 84.2 | -7.1 | 72.5 | 21.7 |
| **2** | -7.6 | -9.4 | 76.2 | -8.1 | 70.3 | -1.7 |
| **3** | -8 | -2.5 | 80.6 | -10.3 | 73.2 | 2.6 |
| **4** | -8 | -8.9 | 77.8 | -17.7 | 70.8 | 7.5 |
| **5** | -6.9 | -8.8 | 88.6 | -14.5 | 73.4 | 25.4 |
| **6** | -7.9 | -7.4 | 82.3 | -9.4 | 72.6 | 8.2 |
| **7** | -11.9 | -8.1 | 81.7 | -9 | 65.3 | 19 |
| **8** | -7.8 | -1.6 | 88 | -13.3 | 73 | 9.7 |
| **9** | -8.3 | -9.9 | 86.1 | -10 | 76 | 6.3 |
| **10** | -7.5 | -10 | 86.2 | -10.7 | 73.7 | 4 |
| **11** | -7.5 | -6.6 | 79.7 | -9.9 | 72.5 | 12.9 |
| **12** | -7.6 | -5.4 | 86.3 | -8.4 | 69.7 | 3.4 |
| **13** | -10.4 | 2.1 | 74.3 | -8.7 | 72.8 | -1.1 |
| **14** | -8 | -9.8 | 76.3 | -5.8 | 72.9 | 2.6 |
| **15** | -6.7 | -10.6 | 84.9 | -8 | 70.9 | 3.7 |
| **16** | -8.3 | -10.5 | 84.9 | -7.3 | 73.5 | 11.1 |
| **17** | -7.4 | -6 | 79.9 | -6.4 | 73.9 | 8.1 |
| **18** | -7.6 | -10.9 | 85.4 | -8.5 | 76 | 9.6 |
| **19** | -7.2 | -7.1 | 85.7 | -8.5 | 73.8 | 5.8 |
| **20** | -7.8 | -9.6 | 88.4 | -5.9 | 74.8 | 5.8 |
| **Mean** | **-7.9** | **-7.4** | **82.9** | **-9.4** | **72.6** | **8.2** |
| **SE** | **0.3** | **0.8** | **1** | **0.7** | **0.5** | **1.7** |

For both SMA and mOFC stimulation, the center of the coil was placed over the corresponding target location (see Figure 1). In SMA blocks, the coil was held tangential to the scalp with the handle pointing at a -100° angle away from the midsagittal line (*i.e.*, in the counter-clockwise direction), resulting in a medio-lateral current flow within the cortex (Figure 1). This coil position was chosen based on a previous experiment showing that it allows the most optimal recruitment of SMA neurons (Arai et al., 2012). In mOFC blocks, the coil was held tangential to the forefront with the handle directed upward and parallel to the midsagittal line (Codol et al., 2020), resulting in a downward current flow at the cortical level.

#### Stimulation intensities

Once the real and the adjusted hotspots were found (see above), we determined the resting motor threshold (rMT) for both locations. The rMT was defined as the lowest stimulation intensity (expressed in percentage of maximal stimulator output (%MSO)) required to evoke MEPs of 50 μV amplitude on 5 out of 10 consecutive trials in the relaxed FDI muscle (Rossini et al., 1994, 2015). The rMTs for the real and the adjusted hotspots were 39.9 ± 1.5 % and 44.6 ± 1.9 %MSO, respectively (see Table 2 for individual rMT values).

**Table 2:** Resting motor threshold (rMT) expressed in percentage of maximal stimulator output (%MSO) obtained for each subject using the real and adjusted hotspots. Subjects for whom the hotspot had to be adjusted (12/20 subjects) are highlighted in black.

| Subject | Real hotspot<br>rMT (%MSO) | Adjusted hotspot<br>rMT (%MSO) |
| --- | --- | --- |
| 1 | 37 | 37 |
| 2 | 31 | 37 |
| 3 | 36 | 45 |
| 4 | 39 | 39 |
| 5 | 45 | 55 |
| 6 | 51 | 59 |
| 7 | 50 | 54 |
| 8 | 40 | 45 |
| 9 | 31 | 31 |
| 10 | 37 | 37 |
| 11 | 42 | 42 |
| 12 | 35 | 44 |
| 13 | 57 | 65 |
| 14 | 41 | 41 |
| 15 | 35 | 35 |
| 16 | 38 | 46 |
| 17 | 43 | 48 |
| 18 | 38 | 42 |
| 19 | 41 | 49 |
| 20 | 39 | 39 |
| Mean | 39.9 | 44.6 |
| SE | 1.5 | 1.9 |

These rMT values were exploited to determine the stimulation intensities to be used for the rest of the experiment. In SMA blocks, the M1 coil was positioned over the adjusted hotspot. Hence, we based on the rMT obtained at this location to define the stimulation intensity for M1; we stimulated M1 at 120 % of this rMT (Derosiere et al., 2017a, 2017b). Conversely, in mOFC blocks, the M1 coil could be easily positioned over the real hotspot and we thus stimulated at 120 % of the rMT obtained for the real hotspot. Finally, CS intensity was set at 120 % of the rMT obtained for the real hotspot, both in SMA and in mOFC blocks (Brown et al., 2019).

#### Inter-stimulation intervals and blocks

As mentioned in the Introduction section, the goal of the present study was fourfold. First, we aimed to test the influence of SMA stimulation on M1 activity with long inter-stimulation intervals, presumed to recruit cortico-subcortico-cortical circuits. To this aim, we exploited intervals of 12 and 15 ms (Neubert et al., 2010). As a second objective, we sought to clarify whether the MEP potentiation reported in studies targeting SMA and M1 with short intervals (Arai et al., 2011, 2012; Green et al., 2018; Rurak et al., 2021) can be taken as a pure measure of cortico-cortical connectivity between these areas or if it could in part reflect the summation of volleys descending from those on motoneurons. To address this issue, we tested the effect of SMA conditioning on MEP amplitudes using a very short inter-stimulation intervals of 1 ms. The rationale here was that a 1 ms interval would be too short for a MEP potentiation to result from the recruitment of cortico-cortical circuits (Tokuno & Nambu, 2000; Aizawa & Tanji, 1994); any potentiation occurring with this interval would instead provide evidence for a summation of neural inputs occurring at the spinal level. We also included other short inter-stimulation intervals of 4, 6 and 8 ms in the experiment to be able to compare the effect obtained on MEP amplitudes when using the 1 ms interval *vs.* when exploiting more classical intervals. Finally, we aimed to test the feasibility of exploiting ppTMS to probe effective connectivity between mOFC and M1 and, relatedly, to determine the influence of mOFC stimulation on motor activity. Given the current lack of data regarding the latter issue, we exploited all of the intervals mentioned above in mOFC blocks too. Note that, contrary to SMA, mOFC does not present corticospinal projections. Hence, the use of a 1 ms interval in mOFC blocks allowed us to verify that any effect of SMA stimulation on MEPs using this interval was specific to SMA. Altogether, the experiment involved 6 inter-stimulation intervals, both in SMA and in mOFC blocks: 1, 4, 6, 8, 12 and 15 ms.

The experiment was divided into 10 blocks of 42 trials (*i.e.*, 5 SMA blocks and 5 mOFC blocks). Each block comprised trials with single-pulse (*i.e.*, TS only) and paired-pulse TMS (*i.e.*, CS+TS with the 6 intervals mentioned above), occurring in a randomized order. As such, within each block, a total of 6 trials was recorded for each of the 7 conditions (*i.e.*, single-pulse, and paired-pulse with 1, 4, 6, 8, 12 and 15 ms), leading to 30 trials per condition over the whole experiment. These high numbers are quite unusual for TMS studies (*i.e.*, which typically involve 10 to 20 MEPs per condition; *e.g.*, see Arai et al., 2012; Beaulieu et al., 2017; Neige et al., 2017; Derosiere et al., 2019). Having a large number of MEPs reduces within-subject variability and may help increase the reliability of the findings (Chang et al., 2016; Beaulieu et al., 2017). Finally, to prevent subjects from anticipating the stimulations, we varied the inter-trial interval, which ranged between 3.6 and 4.4 s (*i.e.*, rectangular distribution centered over 4 s; Rothwell et al., 1999).

### Electromyographic recordings

Electromyography was used to record MEPs in the right FDI muscle. To do so, pairs of surface electrodes (Ag/AgCl, Medicotest, USA) were disposed on the FDI in a belly-tendon montage. A ground electrode was placed on the styloid process of the right ulna. EMG signals were recorded for 800 ms on each trial, starting and ending 400 ms before and after the TS, respectively. The signals were amplified with a gain of 1000, and band-pass and notch-filtered (10–500 Hz and 50 Hz, respectively) online using a dedicated amplifier (Digitimer D360; Digitimer Ltd., Welwyn Garden City, UK). Signals were digitized at a sampling rate of 2000 Hz (CED Power 1401; CED Ltd., Cambridge, UK) and collected using the Signal software (version 6.04; CED Ltd.) for further offline analyses.

### Data analyses

#### EMG data

EMG data were analyzed with custom Signal and R scripts (R Core Team, 2020). Of note, to prevent contamination of the MEP measurements from background muscular activity, participants were reminded to relax during the whole experiment based on the EMG signals, which were continuously screened by the experimenters. In addition, trials in which the root mean square of the EMG signal exceeded 2.5 SD above the mean before stimulation (*i.e.*, - 250 to -50 ms from the pulse) were discarded from the analyses. Besides, to attenuate any effect of MEP variability on our measures, MEPs with an amplitude exceeding 2.5 SD around the mean within a given condition were excluded too. Following this cleaning procedure, we had 84.9 ± 7.55 % and 86.63 ± 5.53 % trials left on average for SMA and mOFC blocks, respectively.

We then extracted the peak-to-peak MEP amplitude for each subject, each condition and each single trial. Trials were subsequently pooled together, by computing the median amplitude for each subject and each condition. The nature of the influence of the SMA/the mOFC over M1 (*i.e.*, facilitatory *vs*. suppressive) was finally quantified by computing a ratio expressing MEPs elicited by the TS in paired-pulse trials relative to MEPs elicited in the TS in single-pulse trials (Lafleur et al., 2016; Derosiere et al., 2020; Koch, 2020; Neige et al., 2021). Following this procedure, one MEP ratio was obtained for each subject and each inter-stimulation interval. MEP ratios above 1 were taken as a marker of a facilitatory influence of the conditioned area on M1, whereas ratios below 1 were considered as reflecting a suppressive effect on M1.

#### MRI data

Individual MRI volumetric data were used to investigate whether the facilitatory/suppressive effect of the SMA/mOFC over M1, demonstrated by dual-site ppTMS at specific, significant intervals, could be related to the cortical volume of these two areas.

The segmentation and parcellation of brain cortical and subcortical regions were performed using FreeSurfer (version 7.2.; http://surfer.nmr.harvard.edu). Following the completion of the pipeline, each segmentation was visually inspected and corrected if necessary. Minor errors in the cranial stripes were identified and rectified before re-running the data through the pipeline. The regions of interest (ROIs) were defined based on the Brainnetome atlas (https://atlas.brainnetome.org/) (Fan et al., 2016), which provides a fine-grained, connectivity-based parcellation of the human brain into 246 regions. Specifically, we focused on the left SMA (region 9 from the Brainnetome Atlas) and the left mOFC (regions 41 and 47) which were projected to the Freesurfer’s parcellation of each individual’s brain. As a control area, the occipital polar cortex (region 204) was also selected. Next, the automated pipeline enabled the generation of cerebral volume measures (in mm³). The volume obtained for each ROI was normalized to the total intracranial volume of the subject to account for individual brain size.

### Statistical analyses

All statistical analyses were performed using the JASP software (JASP Team, version 0.14.1.0; https://jasp-stats.org/).

First, we aimed to identify whether the influence of the CS on MEP amplitudes varied as a function of the inter-stimulation interval (*e.g.*, whether the effect for an interval of 1 vs. 4, 6 and 8 ms differed for SMA blocks; see section *Inter-stimulation intervals and blocks*, above). Data were normally distributed, as evidenced by non-significant results of the Shapiro-Wilk tests. When running repeated-measures (rm)ANOVA, Mauchly’s tests were systematically exploited to check for data sphericity and a Greenhouse-Geiser correction was applied if the sphericity assumption was violated. To address this point, MEP ratios obtained for SMA and mOFC blocks were analyzed using two separate one-way rmANOVAs with INTERVAL (1, 4, 6, 8, 12 and 15 ms) as a within-subject factor. Pre-planned post-hoc analyses were performed on significant interactions after applying a Bonferroni correction for multiple comparisons. Effect sizes were estimated for the main effect of INTERVAL, by calculating partial eta squared (η^2^_p_). In accordance with conventional interpretation partial η^2^_p_, a value of 0.01 is interpreted as indicating a small effect size, a value of 0.06 a medium effect size and a value of 0.14 or more as a large effect size (Lakens, 2013). Second, we sought to determine whether the CS had a significant facilitatory or suppressive effect on MEP amplitude. To this aim, MEP ratios were compared against a constant value of 1 (*i.e.*, reflecting the amplitude obtained in TS only trials) using one-sample t-tests, as usually performed in TMS studies (Arai et al., 2012; Quoilin et al., 2019; Neige et al., 2020; Wang et al., 2020).

To complement the frequentist statistics, we conducted a Bayes factor (BF) analysis, allowing us to quantify statistically the level of evidence for the presence of an effect on MEP ratios. These analyses were performed using the JASP default parameters (*i.e.*, Cauchy prior width of 0.707; van Doorn et al., 2021). BFs (expressed as BF_10_) provided us with a ratio of the likelihood probability of the alternative hypothesis (*i.e.*, H_1_: the probability that data exhibit the effect; Morey & Rouder, 2011) over the null hypothesis (*i.e.*, H_0_: the probability that data do not exhibit an effect of the tested factor). A BF_10_ of 1 reflect an equal probability that H_1_ and H_0_ are correct, whereas a value higher than 1 would reflect a higher probability that H_1_ is correct. In accordance with conventional interpretation of BF values (Jeffreys, 1961), a BF_10_ value ranging between 1 and 3 is interpreted as indicating anecdotal evidence in favor of H_1_, a value between 3 and 10 as indicating substantial evidence for H_1_, a value between 10 and 30 a strong evidence for H_1_, a value between 30 and 100 a very strong evidence for H_1_, and a value above 100 a decisive evidence for H_1_. Conversely, a BF_10_ value between 0.1–0.33 and 0.33–1 indicates substantial and anecdotal evidence for the null hypothesis, respectively.

Lastly, we aimed to determine whether the individual cortical volumes of the SMA and the mOFC could account for their observed facilitatory/suppressive influence on MEP amplitudes. To assess these non-normally distributed data, Spearman’s correlations (nonparametric) were performed between the mean MEP ratio for significant facilitatory/suppressive intervals and the respective normalized cortical volume values of the SMA and the mOFC. Additionally, to ensure that the correlation did not depend on age- and gender-related differences in cortical morphology, Spearman’s partial correlations were performed to confirm our results by considering age and gender as control variables. Finally, evaluate the specificity of these findings, Spearman’s correlations were also performed between the same MEP ratios and the normalized cortical volume of a control occipital polar area, as well as the normalized total gray matter volume.

## Results

### SMA stimulation induced facilitatory and suppressive effects on MEP amplitudes depending on the inter-stimulation interval

Figures 2.A and B illustrate the MEP ratios obtained as a function of the inter-stimulation interval for SMA blocks. Interestingly, the rmANOVA revealed a main effect of the factor INTERVAL on MEP ratios (GG-corrected F_(2.41,45.87)_ = 10.75, p < .0001). The n^2^ for this effect was .361, denoting a large effect size. Further, the BF_10_ was 560807, indicative of a ‘decisive’ evidence in favor of H_1_ (*i.e.*, H_1_: presence of an effect of INTERVAL) over H_0_ (*i.e.*, H_0_: lack of effect of INTERVAL). Post-hoc analyses indeed showed that MEP ratios strongly varied as a function of the inter-stimulation interval: ratios for 6 ms (p < .001; BF_10_ = 114.17), 8 ms (p < .001; BF_10_ = 87.44), 12 ms (p < .001; BF_10_ = 35.29) and 15 ms (p < .001; BF_10_ = 119.87) were significantly higher than at 1 ms. Moreover, ratios for intervals of 8 ms (p < .001; BF_10_ = 12.67) and 12 ms (p = .006; BF_10_ = 8.46) were significantly higher than for an interval of 4 ms. Finally, the ratio at 12 ms was not significantly different from ratios obtained at 6 ms (p > .999; BF_10_ = 0.603) and 8 ms (p > .999; BF_10_ = 0.261) intervals, nor was the ratio at 15 ms when compared to 6 ms and 8 ms intervals (p > .999, BF_10_ = 0.246 and p > .999, BF_10_ = 1.235, respectively). Altogether, these findings indicate that the facilitatory effect of SMA stimulation reported in the literature using classical, short intervals (*i.e.*, 6 to 8 ms; Arai et al., 2011, 2012; Green et al., 2018; Rurak et al., 2021) was significantly stronger than when using a 1 ms interval. Further, the similarity of MEP ratios obtained with intervals of 6-8 ms and of 12-15 ms suggests that the facilitatory effect of SMA stimulation reported for short intervals – supposed to probe cortico-cortical circuits – was also present at longer intervals – probing cortico-subcortico-cortical circuits preferentially (Neubert et al., 2010).

**Figure 2:**
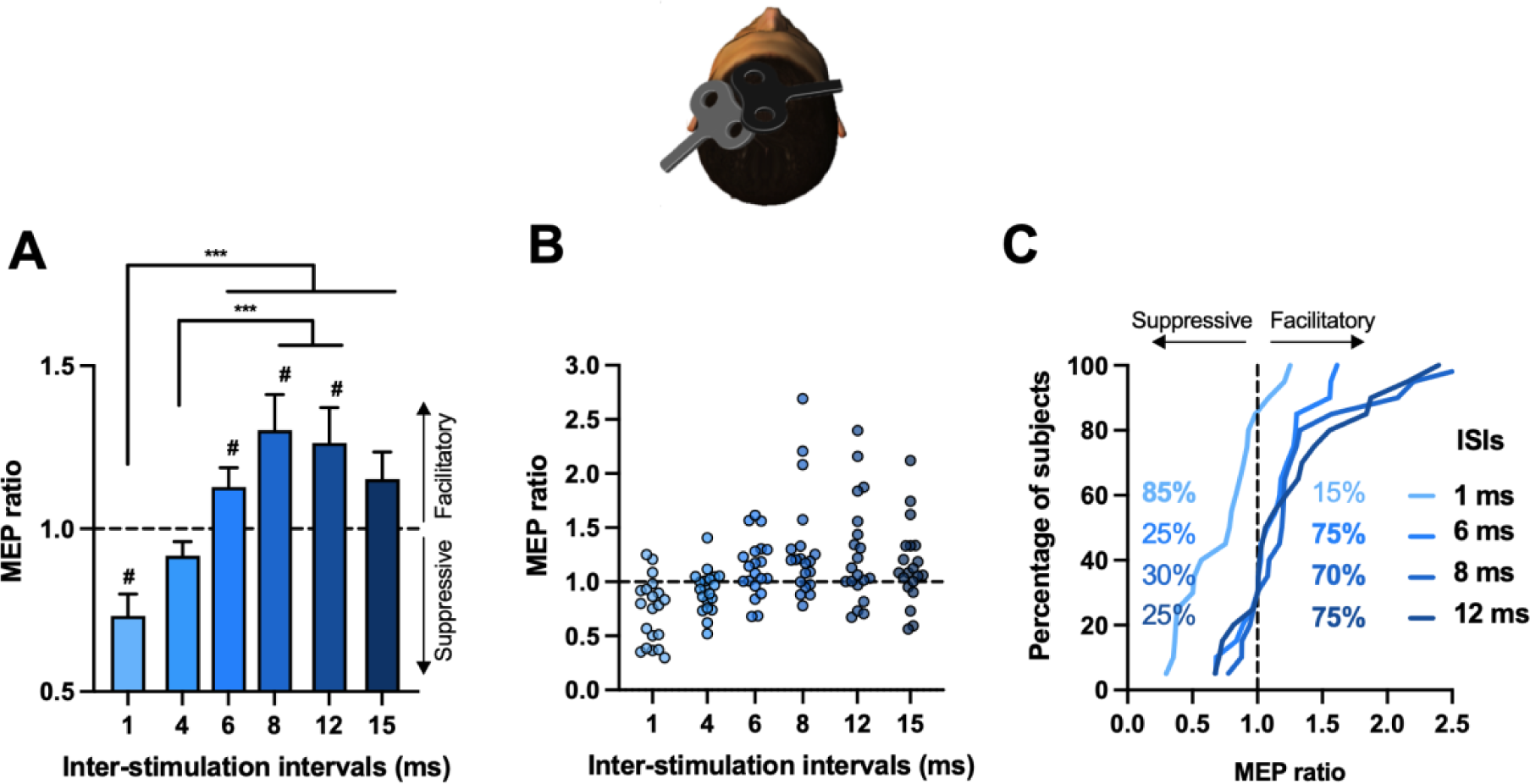
SMA stimulation induced a mix of facilitatory and suppressive effects on MEP amplitudes depending on the inter-stimulation interval. **A.** Group-averaged MEP ratios. Ratios above 1 indicate a facilitatory influence of SMA on M1, whereas ratios below 1 reflect a suppressive effect on M1. *** indicates significant differences between inter-stimulation intervals at p < .001). # indicates a significant difference of the ratio with respect to 1. Error bars represent 1 SEM. **B.** Individual MEP ratios for each inter-stimulation intervals. **C.** Cumulative percentage of subjects for inter-stimulation intervals at which MEP ratios were significantly different than 1. 75, 70 and 75 % of subjects presented a ratio above 1 at 6, 8 and 12 ms, respectively (dark blue traces), while 85 % of subjects presented a ratio below 1 when using the 1 ms interval (light blue trace).

As mentioned above, to directly test whether the CS had a significant facilitatory or suppressive effect on MEP amplitudes, MEP ratios were compared to a constant value of 1 (i.e., representing the mean amplitude of test MEPs) using one-sample t-tests. Interestingly, this analysis confirmed the presence of a significant facilitatory influence of SMA stimulation on MEP amplitudes (*i.e.*, MEP ratios > 1) for classical, short intervals of 6 ms (*t*_19_ = 2.161, p = .044; BF_10_ = 1.55) and 8 ms (*t*_19_ = 2.766, p = .012; BF_10_ = 4.32), in accordance with the literature. Most importantly, a similar facilitatory effect was found for 12 ms (*t*_19_ = 2.435, p = .025; BF_10_ = 2.42). Of note, these effects did not reach statistical significance anymore if corrected for multiple testing by a conservative Bonferroni correction (*i.e*., p-value threshold = .05 / 6 = .008). However, as highlighted above, BF analyses revealed a moderate to strong evidence for the presence of a facilitatory effect at these three inter-stimulation intervals (average BF_10_ = 2.76 ± 0.81), indicating that the data were 2.76 times more likely to show facilitation than no difference from the constant value of 1. As such, 75 %, 70 % and again 75 % of subjects had a MEP ratio above 1 at 6, 8 and 12 ms, respectively, indicating that SMA stimulation potentiated MEP amplitudes for most of the subjects at these intervals (see Figure 2.C). Finally, another interesting finding revealed by the t-tests was the presence of a significant suppressive influence of SMA stimulation on MEP amplitudes for the 1 ms interval (*t*_19_ = 4.034, p < .001; BF_10_ = 49.39). In fact, 85 % of subjects presented a MEP ratio below 1 when using the 1 ms interval, indicating that SMA stimulation had a suppressive effect for a very large proportion of subjects. Other t-tests failed to achieve the level of statistical significance (p-values were of .081 and .070 for 4 and 15 ms).

### mOFC stimulation induced a moderate, interval-dependent suppressive effect on MEP amplitudes

Figures 3.A and B. illustrate the MEP ratios obtained as a function of the inter-stimulation interval for mOFC blocks. The rmANOVA performed on MEP ratio did not indicate a significant main effect of INTERVAL (F_(1,19)_ = 1.148, P =.340, n^2^ = .057; BF_10_ = 0.14).

**Figure 3:**
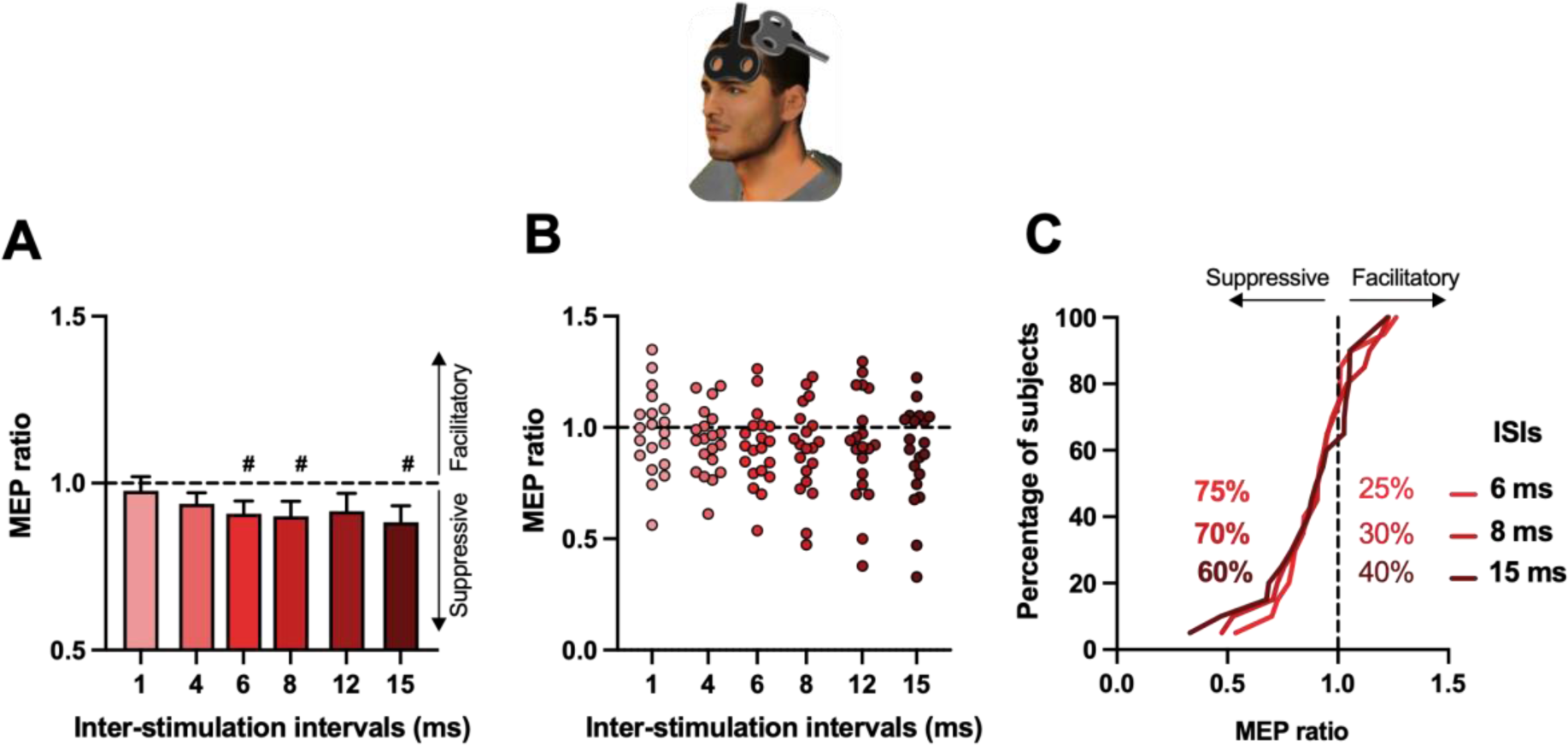
mOFC stimulation induced a moderate suppressive effect on MEP amplitudes at some inter-stimulation intervals. **A.** Group-averaged MEP ratios. Ratios below 1 reflect a suppressive effect on M1. # indicates a significant difference of the ratio with respect to 1. Error bars represent 1 SEM. **B.** Individual MEP ratios for each inter-stimulation intervals. **C.** Cumulative percentage of subjects for inter-stimulation intervals at which MEP ratios were significantly different than 1. 75, 70 and 60 % of subjects presented a ratio below 1 at 6, 8 and 15 ms, respectively.

Despite this lack of a main effect of INTERVAL, we tested whether the CS had a significant facilitatory or suppressive effect on MEP amplitudes at specific intervals (*i.e.*, by comparing ratios to a constant value of 1 with t-tests). Interestingly, this analysis revealed a significant suppressive influence of mOFC stimulation on MEP amplitudes (*i.e.*, MEP ratios < 1) for short intervals of 6 ms (*t*_19_ = -2.425, p = .025; BF_10_ = 2.38) and 8 ms (*t*_19_ = -2.201, p = .040; BF_10_ = 1.65) as well as for a longer interval of 15 ms (*t*_19_ = -2.370; p = .029, BF_10_ = 2.17). Although the BFs for these effects provided anecdotal evidence in favor of H_1_, a total of 75 %, 70 %, and 60 % of subjects presented a MEP ratio greater than 1 at intervals of 6, 8 and 15 ms, respectively. This indicates that mOFC stimulation decreased MEP amplitudes for a large proportion of subjects at these intervals (Figure 3.C). Interestingly, other t-tests did not reach the level of statistical significance (all p-values ranged from .077 to .595), including for the ratio obtained with the 1 ms interval (*t*_19_ = -0.541, p = .595; BF_10_ = 0.265). The latter observation suggests that the suppressive effect observed with this interval was specific to SMA conditioning.

### The facilitatory influence of SMA stimulation on M1 depends on SMA volume

Figures 4.A and B. present the ROIs and the correlations between MEP ratios for significant facilitatory intervals (mean MEP ratios at 6, 8, and 12 ms) and the normalized cortical volumes for the SMA and the mOFC. Spearman’s rank-order correlation analysis revealed a positive association between the facilitatory effect of the SMA over M1 and greater SMA cortical volume (Spearman’s Rho = .682, p = .002). Importantly, this correlation was also present when considering age and gender as control variables in a Spearman partial correlation analyses (Spearman’s Rho = .612, p = .008). Further, this association was specific to the SMA cortical volume, as no significant correlations were observed between the mean MEP ratio and the control occipital polar cortex (Spearman’s Rho = .051; p = .837) or total gray matter volume (Spearman’s Rho = .256; p = .289). In other words, individuals with a larger SMA volume demonstrated a more pronounced facilitatory effect of SMA over M1 at 6, 8, and 12 ms intervals.

**Figure 4:**
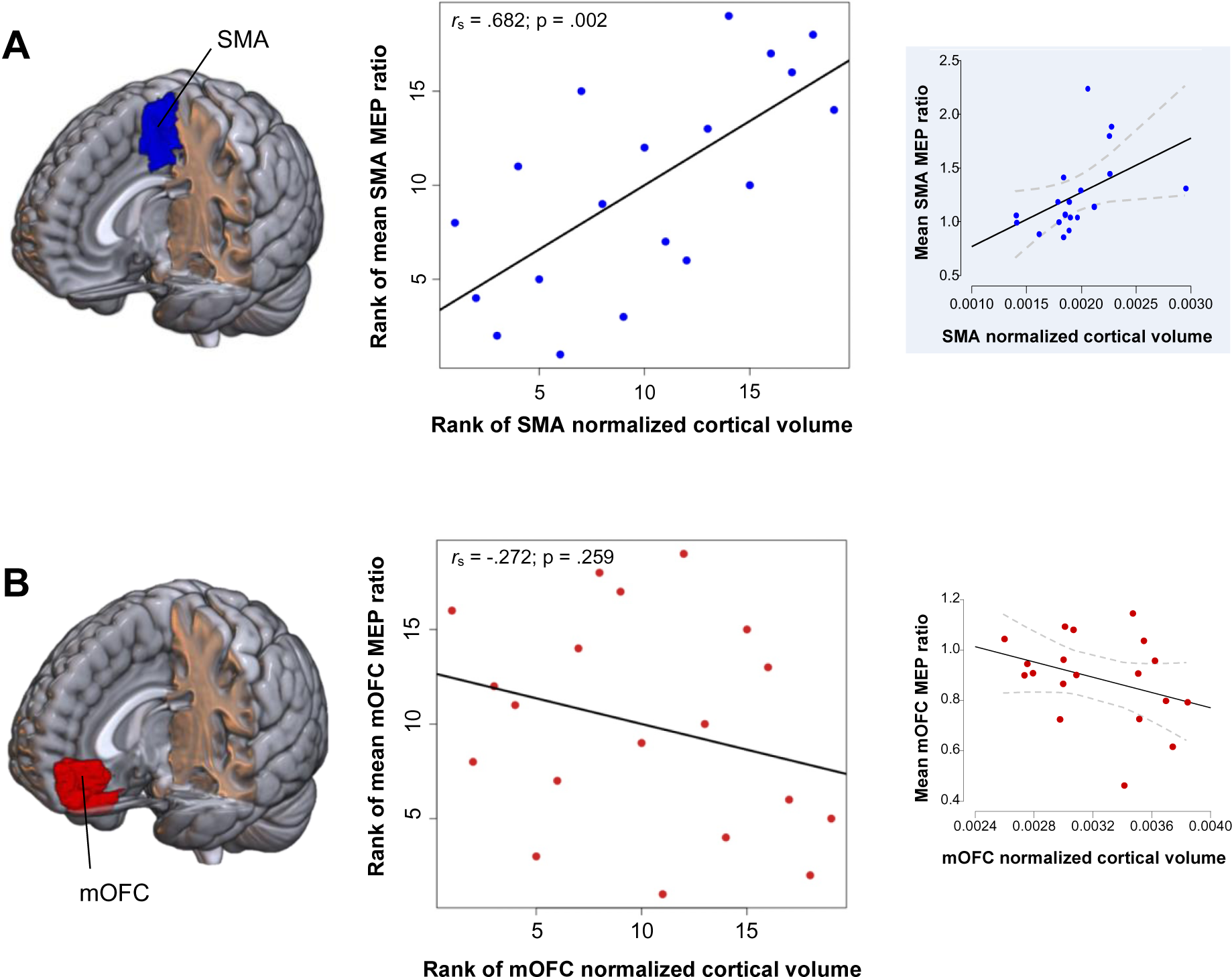
The cortical volume of the SMA (but not the mOFC) contributes to the facilitatory influence on MEP amplitude observed in the dual-site ppTMS data. **A.** 3D representation of the SMA, area 9 of the Brainnetome Atlas. Significant Spearman’s correlation between the mean MEP ratios of the SMA (6, 8 and 12 ms intervals) and the SMA normalized cortical volume. The Pearson correlation between non-ranked data is visualized in the blue window, on the right side of the Figure. **B.** 3D representation of the mOFC, encompassing areas 41 and 47 of the Brainnetome Atlas. Non-significant Spearman’s correlation between mean MEP ratios of the mOFC (6, 8 and 15 ms intervals) and normalized cortical volume of the mOFC. The Pearson correlation between non-ranked data is visualized in the red window. The 3D images were generated using MRIcroGL software (v1.0.20220720). r_s_: Spearman’s Rho.

In contrast, Spearman’s correlations indicated that the inhibitory effect of mOFC over M1 (mean MEP ratios obtained at 6, 8 and 15 ms) did not show a significant association with mOFC cortical volume (Spearman’s Rho = -0.221; p = .356; results of the partial correlation considering age and gender as control variables: Rho = -0.213; p = .412).

## Discussion

Over the past two decades, dual-site ppTMS has been widely used in humans, with studies probing the causal influence of multiple frontal and parietal areas on M1. However, several important questions currently remain unanswered, notably regarding the application of this approach to key areas of the medial frontal cortex – including SMA and mOFC. The present study directly addressed three of these issues. First, we aimed to provide insight into the influence of SMA stimulation on M1 activity with long inter-stimulation intervals (12 to 15 ms), which are thought to recruit cortico-subcortico-cortical circuits (Neubert et al., 2010). Our data reveal that SMA stimulation significantly potentiates MEP amplitudes with a 12 ms interval, indicating the recruitment of circuits that bear a facilitatory influence on M1. Second, we sought to clarify whether the MEP potentiation reported in studies targeting SMA and M1 with short intervals (6 to 8 ms) can be taken as a pure measure of cortico-cortical connectivity between these areas or whether it might in part reflect the summation of descending volleys on motoneurons at the spinal level. Here, we were able to replicate the MEP potentiation previously observed for such intervals. More importantly, our data show that this facilitation does not occur when using a very short interval of 1 ms, assumed to recruit spinal circuits. Rather, we found a suppressive influence of SMA stimulation on MEP amplitudes at this interval. Finally, we tested the feasibility of exploiting ppTMS to probe effective connectivity between mOFC and M1 and, relatedly, we determined the influence of mOFC stimulation on motor activity. We found that mOFC stimulation induced a moderate suppressive effect on MEP amplitudes with both short and long inter-stimulation intervals. Interestingly, mOFC stimulation did not alter MEP amplitudes with the 1 ms interval, suggesting that the suppressive effect observed with this interval was specific to SMA conditioning.

As mentioned above, SMA stimulation induced a significant potentiation of MEP amplitudes with the 12 ms interval. This finding is not trivial because SMA projects to M1 through multiple cortico-subcortico-cortical circuits (Nachev et al., 2008; Accolla et al., 2016; Oswal et al., 2021), some of which exert a net facilitatory influence on motor activity (*e.g.*, the direct pathway of the basal ganglia) and others of which play a suppressive role (*e.g.*, the indirect and hyperdirect pathways). One possibility is that SMA stimulation preferentially recruits the direct pathway of the basal ganglia. In this pathway, areas of the frontal cortex (including SMA) rely on their projections to the striatum to inhibit the internal segment of the globus pallidus, which in turn suppresses neural activity in the subthalamic nucleus (Alexander & Crutcher, 1990; Aron et al., 2007; Calabresi et al., 2014; Niranjan et al., 2018). Since the latter structure exerts a suppressive influence on the motor system (Frank, 2006; Aron et al., 2016; Quartarone et al., 2020), the recruitment of this whole circuit ultimately leads to a disinhibition of M1, putatively explaining the MEP potentiation observed at 12 ms interval. Interestingly, the potentiation did not reach statistical significance when a 15 ms interval was used, suggesting that the facilitatory effect uncovered here is interval-dependent, with an optimal temporal window of about 12 ms. The latter observation is relevant for future ppTMS studies aimed at investigating these circuits.

Of note, we were able to replicate the MEP potentiation previously observed for intervals of 6 and 8 ms (Arai et al., 2011, 2012; Green et al., 2018; Rurak et al., 2021). More importantly, our data show that this facilitatory effect does not arise when using a very short interval of 1 ms, presumed to recruit spinal circuits. Therefore, it is reasonable to assume that the MEP potentiation reported in ppTMS studies using intervals of 6 to 8 ms does not result from the summation of excitatory volleys descending from SMA to motoneurons. In fact, we found a suppressive influence of SMA stimulation on MEP amplitudes when using the 1 ms interval, which was present in up to 85 % of the subjects. Although this effect was quite unexpected, a closer look at the literature reveals that a similar reduction in MEP amplitudes could also be observed when the primary somatosensory cortex ipsilateral to the target M1 was stimulating with a 1 ms interval (Brown et al., 2019). Interestingly, the primary somatosensory cortex also has pyramidal cells that project to the spine. Still, this does not explain the reduction of MEP amplitudes. One possible explanation is that corticospinal cells do not exclusively synapse with motoneurons through direct, excitatory connections, which in fact represent only a minority of cortico-motoneuronal connections (Lemon, 2008). Rather, a large proportion of cells connect to motoneurons through complex inhibitory circuits. The SMA, for instance, innervates a specific set of spinal interneurons (Cheney et al., 2004). One possibility is that SMA stimulation led to the recruitment of inhibitory interneurons that reduced motoneurons excitability and ultimately reduced the amplitude of MEPs elicited by M1 stimulation. As highlighted above, our data show that the stimulation of mOFC – which does not contain pyramidal cells – did not reduce MEP amplitudes when the 1 ms interval was used. Similarly, we showed in two previous studies that conditioning the contralateral M1 – which has pyramidal cells that mostly project to the other side of the spine – with a 1 ms interval does not alter MEP amplitudes neither (Grandjean et al., 2018; Vassiliadis et al., 2018). Taken together, these findings indicate that the suppressive effect observed with a 1 ms interval occurs specifically when stimulating areas that present pyramidal cells *and* project to the same side of the spine as the targeted M1.

Another central aim of the current study was to test the feasibility of using dual-site ppTMS to probe effective connectivity between mOFC and M1, and to determine the influence of mOFC stimulation on motor activity. In fact, while former investigations on caudal areas of the medial frontal cortex (*i.e.*, SMA and preSMA) generally reported a facilitatory influence on M1, ppTMS studies on more rostral areas of the frontal lobe, such as the dorsolateral PFC, revealed the operation of suppressive circuits (Wang et al., 2020). Here, we found that mOFC stimulation moderately reduced MEP amplitudes at specific short and long inter-stimulation intervals. This finding suggests the existence of two frontal regions with opposite influences on the motor system at rest, with more caudal areas – *e.g.*, SMA and preSMA – bearing a facilitatory influence preferentially, and more rostral ones – *e.g.*, mOFC and the dorsolateral PFC – exerting a suppressive influence. The opposing influence of these regions on M1 may allow them to implement distinct functional roles in motor behavior.

Previous studies utilizing dual-site TMS have consistently observed variations in effective connectivity between cortical areas and M1, as measured by intra-sample variations in MEP ratios (Ferbert et al., 1992; Rurak et al., 2021). However, the underlying neurophysiological mechanisms responsible for this inter-individual variability have remained elusive. Our structural MRI analyses offer a compelling explanation for this crucial issue, as they reveal that individuals with larger SMA volumes demonstrate a stronger facilitatory influence of SMA stimulation on M1 activity. Existing evidence also supports a link between M1 cortical volume and inter-individual differences in M1 excitability, measured through intra-sample variations in MEP amplitudes (Rosso et al., 2017; Dayan et al., 2018). These findings collectively suggest that greater cortical tissue volume may be associated with an increased number of cells projecting to other neural structures. Consequently, when employing a dual-site ppTMS approach to stimulate the SMA, a broader population of cortico-cortical cells projecting to M1 may be recruited. In the same line, stimulating M1 with a single-pulse TMS approach may activate a wider population of corticospinal neurons projecting to motoneurons. Therefore, our results suggest the presence of a common source of inter-individual variability in connectivity across different circuits of the nervous system.

However, cortical volume alone does not account for all aspects of this inter-individual variability. Notably, we did not find any association between mOFC cortical volume and its suppressive influence on M1 activity. In fact, it is likely that the suppressive influence of mOFC stimulation on M1 occurs indirectly through polysynaptic connections (Derosiere & Duque, 2020), thereby reducing the contribution of mOFC cortical volume to inter-individual differences in effective connectivity. Future studies should explore the potential contribution of other anatomical features to inter-individual variability in effective connectivity (Neubert et al., 2010).

While the current findings offer promising prospects for future ppTMS studies, it is essential to address some methodological considerations. The suppressive effects of mOFC stimulation on MEP amplitudes (*i.e.*, with 6, 8, and 15 ms intervals) exhibited substantial inter-individual variability, resulting in marginal group-effects level. A potential factor contributing to this inter-individual variability is the presumed difficulty in reaching the mOFC with the magnetic field (Dancy et al., 2023). One way to enhance the recruitment of mOFC neurons would be to adjust the intensity of the conditioning stimulation according to the individual scalp-to-cortex distance (Stokes, 2005; Stokes et al., 2007, 2013) as previously done in repetitive TMS studies (Hanlon et al., 2017; Kearney-Ramos et al., 2018). Such an adjustment could result to a 10–30 % increase in intensity given the greater scalp-to-cortex distance for most prefrontal regions relative to M1 (Kähkönen et al., 2004). Nevertheless, this issue warrants further studies, as the application of a high intensity stimulation to this orbitofrontal location may be particularly uncomfortable for the subjects. A second, critical aspect to consider is the coil placement. This is especially true when stimulating SMA due to its spatial proximity with M1. Here, we used a neuronavigation system to target the MNI coordinates of the SMA based on individual MRI images. In the majority of the subjects (n = 12 / 20), we had to slightly adjust the M1 stimulation site (*i.e.*, the hotspot) and its corresponding rMT. To the best of our knowledge, this type of adjustment has never been reported before, despite the existence of several ppTMS studies on areas lying close to M1. For the sake of transparency, we believe that future studies should systematically report any adjustment of coil locations.

Overall, the current study shows that SMA and mOFC conditioning exerts interval- and region-specific facilitatory and suppressive influences on motor activity. Our findings pave the way for both fundamental and clinical investigations aimed at understanding the causal role of these areas in the modulation of motor activity, as may occur in motor planning, decision-making, and inhibitory control, in which SMA and mOFC play a central role.

## Acknowledgements

This work was supported by grants from the Belgian National Funds for Scientific Research (FNRS: MIS F.4512.14) obtained by GD (FNRS: 1B134.18) and from the french ‘Fondation Thérèse & René Planiol’ (Mobility grant) and Wallonie-Bruxelles International (Excellence grant WBI.IN) obtained by CN. P.V. was supported by the Fund for Research training in Industry and Agriculture (FRIA/FNRS; FC29690) and Wallonie-Bruxelles International.

## Data Availability

All datasets will be freely available on the Open Science Framework repository upon publication at https://osf.io/up45j/

## References

Accolla EA, Herrojo Ruiz M, Horn A, Schneider GH, Schmitz-Hübsch T, Draganski B & Kühn AA (2016). Brain networks modulated by subthalamic nucleus deep brain stimulation. Brain 139, 2503–2515.

Aizawa H & Tanji J (1994). Corticocortical and thalamocortical responses of neurons in the monkey primary motor cortex and their relation to a trained motor task. Journal of neurophysiology 71, 550–560.

Alamia A, Zénon A, VanRullen R, Duque J & Derosiere G (2019). Implicit visual cues tune oscillatory motor activity during decision-making. NeuroImage 186, 424–436.

Alexander GE & Crutcher MD (1990). Functional architecture of basal ganglia circuits: neural substrates of parallel processing. Trends in neurosciences 13, 266–271.

Allart E, Devanne H & Delval A (2019). Contribution of transcranial magnetic stimulation in assessing parietofrontal connectivity during gesture production in healthy individuals and brain-injured patients. Neurophysiologie Clinique 49, 115–123.

Amodio DM & Frith CD (2006). Meeting of minds: the medial frontal cortex and social cognition. Nature reviews Neuroscience 7, 268–277.

Arai N, Lu MK, Ugawa Y & Ziemann U (2012). Effective connectivity between human supplementary motor area and primary motor cortex: A paired-coil tms study. Experimental Brain Research 220, 79–87.

Arai N, Muller-Dahlhaus F, Murakami T, Bliem B, Lu MK, Ugawa Y & Ziemann U (2011). State-dependent and timing-dependent bidirectional associative plasticity in the human sma-m1 network. Journal of Neuroscience 31, 15376–15383.

Aron AR, Durston S, Eagle DM, Logan GD, Stinear CM & Stuphorn V (2007). Converging evidence for a fronto-basal-ganglia network for inhibitory control of action and cognition. Journal of Neuroscience 27, 11860–11864.

Aron AR, Herz DM, Brown P, Forstmann BU & Zaghloul K (2016). Frontosubthalamic Circuits for Control of Action and Cognition. The Journal of neuroscience : the official journal of the Society for Neuroscience 36, 11489–11495.

Beaulieu LD, Flamand VH, Massé-Alarie H & Schneider C (2017). Reliability and minimal detectable change of transcranial magnetic stimulation outcomes in healthy adults: A systematic review. Brain Stimulation 10, 196–213.

Bestmann S & Duque J (2016). Transcranial magnetic stimulation: Decomposing the processes underlying action preparation. Neuroscientist 22, 392–405.

Boy F, Husain M, Singh KD & Sumner P (2010). Supplementary motor area activations in unconscious inhibition of voluntary action. Experimental brain research 206, 441– 448.

Brown MJN, Goldenkoff ER, Chen R, Gunraj C & Vesia M (2019). Using dual-site transcranial magnetic stimulation to probe connectivity between the dorsolateral prefrontal cortex and ipsilateral primary motor cortex in humans. Brain Sciences; DOI: 10.3390/brainsci9080177.

Calabresi P, Picconi B, Tozzi A, Ghiglieri V & Di Filippo M (2014). Direct and indirect pathways of basal ganglia: a critical reappraisal. Nature Neuroscience 2014 17:8 17, 1022–1030.

Carlsen AN, Eagles JS & MacKinnon CD (2015). Transcranial direct current stimulation over the supplementary motor area modulates the preparatory activation level in the human motor system. Behavioural brain research 279, 68–75.

Chang WH, Fried PJ, Saxena S, Jannati A, Gomes-Osman J, Kim YH & Pascual-Leone A (2016). Optimal number of pulses as outcome measures of neuronavigated transcranial magnetic stimulation. Clinical neurophysiology : official journal of the International Federation of Clinical Neurophysiology 127, 2892–2897.

Cheney PD, Belhaj-Saïf A & Boudrias M-H (2004). Principles of corticospinal system organization and function. In Clinical Neurophysiology of Motor Neuron Diseases, ed. Eisen A, Handbook of Clinical Neurophysiology, pp. 59–96. Elsevier. Available at: https://www.sciencedirect.com/science/article/pii/S1567423104040043.

Codol O, Galea JM, Jalali R & Holland PJ (2020). Reward-driven enhancements in motor control are robust to TMS manipulation. Experimental Brain Research 238, 1781– 1793.

Dancy MM, Caulfield KA, George MS & Li X (2023). Electric field dosing requires a higher stimulation intensity for medial prefrontal TMS compared with lateral prefrontal TMS. Brain Stimulation 16, 386.

Davare M, Lemon R & Olivier E (2008). Selective modulation of interactions between ventral premotor cortex and primary motor cortex during precision grasping in humans. The Journal of physiology 586, 2735–2742.

Dayan E, López-Alonso V, Liew S-L & Cohen LG (2018). Distributed cortical structural properties contribute to motor cortical excitability and inhibition. Brain Struct Funct 223, 3801–3812.

Derosière G, Billot M, Ward ET & Perrey S (2015). Adaptations of motor neural structures’ activity to lapses in attention. Cerebral Cortex 25, 66–74.

Derosiere G & Duque J (2020). Tuning the Corticospinal System: How Distributed Brain Circuits Shape Human Actions. The Neuroscientist 26, 359–379.

Derosiere G, Klein P-A, Nozaradan S, Zénon A, Mouraux A & Duque J (2018). Visuomotor Correlates of Conflict Expectation in the Context of Motor Decisions. J Neurosci 38, 9486–9504.

Derosiere G, Thura D, Cisek P & Duque J (2019). Motor cortex disruption delays motor processes but not deliberation about action choices. Journal of neurophysiology 122, 1566–1577.

Derosiere G, Vassiliadis P, Demaret S, Zénon A & Duque J (2017a). Learning stage-dependent effect of M1 disruption on value-based motor decisions. Neuroimage 162, 173–185.

Derosiere G, Vassiliadis P & Duque J (2020). Advanced TMS approaches to probe corticospinal excitability during action preparation. NeuroImage 213, 116746.

Derosiere G, Zénon A, Alamia A & Duque J (2017b). Primary motor cortex contributes to the implementation of implicit value-based rules during motor decisions. Neuroimage 146, 1115–1127.

Di Lazzaro V, Rothwell J & Capogna M (2018). Noninvasive Stimulation of the Human Brain: Activation of Multiple Cortical Circuits. Neuroscientist 24, 246–260.

van Doorn J et al. (2021). The JASP guidelines for conducting and reporting a Bayesian analysis. Psychon Bull Rev 28, 813–826.

Dum RP & Strick PL (1991). The origin of corticospinal projections from the premotor areas in the frontal lobe. The Journal of neuroscience : the official journal of the Society for Neuroscience 11, 667–689.

Dum RP & Strick PL (1996). Spinal cord terminations of the medial wall motor areas in macaque monkeys. The Journal of neuroscience : the official journal of the Society for Neuroscience 16, 6513–6525.

Entakli J, Bonnard M, Chen S, Berton E & De Graaf JB (2014). TMS reveals a direct influence of spinal projections from human SMAp on precise force production. European Journal of Neuroscience 39, 132–140.

Fan L, Li H, Zhuo J, Zhang Y, Wang J, Chen L, Yang Z, Chu C, Xie S, Laird AR, Fox PT, Eickhoff SB, Yu C & Jiang T (2016). The Human Brainnetome Atlas: A New Brain Atlas Based on Connectional Architecture. Cerebral Cortex 26, 3508–3526.

Fellows LK (2007). The role of orbitofrontal cortex in decision making. In Annals of the New York Academy of Sciences, pp. 421–430. Blackwell Publishing Inc.

Ferbert A, Priori A, Rothwell JC, Day BL, Colebatch JG & Marsden CD (1992). Interhemispheric inhibition of the human motor cortex. The Journal of physiology 453, 525–546.

Frank MJ (2006). Hold your horses: a dynamic computational role for the subthalamic nucleus in decision making. Neural networks : the official journal of the International Neural Network Society 19, 1120–1136.

Geers L, Pesenti M, Derosiere G, Duque J, Dricot L & Andres M (2021). Role of the fronto-parietal cortex in prospective action judgments. Sci Rep 11, 7454.

Grandjean J, Derosiere G, Vassiliadis P, Quemener L, Wilde Y de & Duque J (2018). Towards assessing corticospinal excitability bilaterally: Validation of a double-coil TMS method. Journal of Neuroscience Methods 293, 162–168.

Green PE, Ridding MC, Hill KD, Semmler JG, Drummond PD & Vallence AM (2018). Supplementary motor area—primary motor cortex facilitation in younger but not older adults. Neurobiology of Aging 64, 85–91.

Hanlon CA, Dowdle LT, Correia B, Mithoefer O, Kearney-Ramos T, Lench D, Griffin M, Anton RF & George MS (2017). Left frontal pole theta burst stimulation decreases orbitofrontal and insula activity in cocaine users and alcohol users. Drug and Alcohol Dependence 178, 310–317.

Hu S & Li C-SR (2012). Neural Processes of Preparatory Control for Stop Signal Inhibition. Human Brain Mapping 33, 2785–2796.

Jeffreys H (1961). The Theory of Probability.

Kähkönen S, Wilenius J, Komssi S & Ilmoniemi RJ (2004). Distinct differences in cortical reactivity of motor and prefrontal cortices to magnetic stimulation. Clinical Neurophysiology 115, 583–588.

Kearney-Ramos TE, Dowdle LT, Lench DH, Mithoefer OJ, Devries WH, George MS, Anton RF & Hanlon CA (2018). Transdiagnostic Effects of Ventromedial Prefrontal Cortex Transcranial Magnetic Stimulation on Cue Reactivity. Biological Psychiatry: Cognitive Neuroscience and Neuroimaging 3, 599–609.

Klein-Flügge MC, Kennerley SW, Friston K & Bestmann S (2016). Neural Signatures of Value Comparison in Human Cingulate Cortex during Decisions Requiring an Effort-Reward Trade-off. The Journal of neuroscience : the official journal of the Society for Neuroscience 36, 10002–10015.

Klöppel S, Bäumer T, Kroeger J, Koch MA, Büchel C, Münchau A & Siebner HR (2008). The cortical motor threshold reflects microstructural properties of cerebral white matter. NeuroImage 40, 1782–1791.

Koch G (2020). Cortico-cortical connectivity: the road from basic neurophysiological interactions to therapeutic applications. Experimental Brain Research 238, 1677– 1684.

Koch G & Rothwell JC (2009). TMS investigations into the task-dependent functional interplay between human posterior parietal and motor cortex. Behavioural Brain Research 202, 147–152.

Koch G, Ruge D, Cheeran B, Fernandez Del Olmo M, Pecchioli C, Marconi B, Versace V, Lo Gerfo E, Torriero S, Oliveri M, Caltagirone C & Rothwell JC (2009). TMS activation of interhemispheric pathways between the posterior parietal cortex and the contralateral motor cortex. The Journal of physiology 587, 4281–4292.

Koch G, Schneider S, Baumer T, Franca M, Munchau A, Cheeran B, Bhatia KP & Rothwell JC (2006). Interhemispheric inhibition of the dorsal premotor-motor pathway is reduced in writer’s cramp dystonia. MOVEMENT DISORDERS 21, S388.

Lafleur L-P, Tremblay S, Whittingstall K & Lepage J-F (2016). Assessment of Effective Connectivity and Plasticity With Dual-Coil Transcranial Magnetic Stimulation. Brain Stimulation 9, 347–355.

Lakens D (2013). Calculating and reporting effect sizes to facilitate cumulative science: A practical primer for t-tests and ANOVAs. Frontiers in Psychology 4, 1–12.

Lebon F, Lotze M, Stinear CM & Byblow WD (2012). Task-Dependent Interaction between Parietal and Contralateral Primary Motor Cortex during Explicit versus Implicit Motor Imagery ed. Wenderoth N. PLoS ONE 7, e37850.

Lemon RN (2008). Descending pathways in motor control. Annual review of neuroscience 31, 195–218.

List J, Kübke JC, Lindenberg R, Külzow N, Kerti L, Witte V & Flöel A (2013). Relationship between excitability, plasticity and thickness of the motor cortex in older adults. Neuroimage 83, 809–816.

Loh MK & Rosenkranz JA (2022). The medial orbitofrontal cortex governs reward-related circuits in an age-dependent manner. Cereb Cortex 33, 1913–1924.

Luppino G, Matelli M, Camarda R & Rizzolatti G (1993). Corticocortical connections of area F3 (SMA-proper) and area F6 (pre-SMA) in the macaque monkey. The Journal of comparative neurology 338, 114–140.

Makoshi Z, Kroliczak G & van Donkelaar P (2011). Human supplementary motor area contribution to predictive motor planning. Journal of Motor Behavior 43, 303–309.

Mars RB, Klein MC, Neubert F-X, Olivier E, Buch ER, Boorman ED & Rushworth MFS (2009). Short-latency influence of medial frontal cortex on primary motor cortex during action selection under conflict. The Journal of neuroscience : the official journal of the Society for Neuroscience 29, 6926–6931.

Morey RD & Rouder JN (2011). Bayes Factor Approaches for Testing Interval Null Hypotheses. Psychological Methods 16, 406–419.

Muakkassa KF & Strick PL (1979). Frontal lobe inputs to primate motor cortex: evidence for four somatotopically organized “premotor” areas. Brain research 177, 176–182.

Nachev P, Kennard C & Husain M (2008). Functional role of the supplementary and pre- supplementary motor areas. Nature Reviews Neuroscience 9, 856–869.

Neige C, Brun C, Gagné M, Bouyer LJ & Mercier C (2020). Do nociceptive stimulation intensity and temporal predictability influence pain-induced corticospinal excitability modulation? NeuroImage116883.

Neige C, Ciechelski V & Lebon F (2022). The recruitment of indirect waves within primary motor cortex during motor imagery: A directional transcranial magnetic stimulation study. Eur J Neurosci; DOI: 10.1111/ejn.15843.

Neige C, Massé-Alarie H, Gagné M, Bouyer LJ & Mercier C (2017). Modulation of corticospinal output in agonist and antagonist proximal arm muscles during motor preparation ed. Tremblay F. PLOS ONE 12, e0188801.

Neige C, Massé-Alarie H & Mercier C (2018). Stimulating the Healthy Brain to Investigate Neural Correlates of Motor Preparation: A Systematic Review. Neural Plasticity 2018, 1–14.

Neige C, Rannaud Monany D & Lebon F (2021). Exploring cortico-cortical interactions during action preparation by means of dual-coil transcranial magnetic stimulation: A systematic review. Neuroscience and Biobehavioral Reviews 128, 678–692.

Neubert FX, Mars RB, Buch ER, Olivier E & Rushworth MF (2010). Cortical and subcortical interactions during action reprogramming and their related white matter pathways. Proceedings of the National Academy of Sciences of the United States of America 107, 13240–13245.

Niranjan A, Lunsford LD & Richardson RM (2018). Current Concepts in Movement Disorder Management. *Prog Neurol Surg Basel*, Karger 33, 50–61.

Oswal A, Cao C, Yeh CH, Neumann WJ, Gratwicke J, Akram H, Horn A, Li D, Zhan S, Zhang C, Wang Q, Zrinzo L, Foltynie T, Limousin P, Bogacz R, Sun B, Husain M, Brown P & Litvak V (2021). Neural signatures of hyperdirect pathway activity in Parkinson’s disease. Nature Communications 2021 12:1 12, 1–14.

Pierre Vassiliadis, Elena Beanato, Traian Popa, Fabienne Windel, Takuya Morishita, Esra Neufeld, Julie Duque, Gerard Derosiere, Maximilian J. Wessel, & Friedhelm C. Hummel (2023). Non-invasive stimulation of the human striatum disrupts reinforcement learning of motor skills. bioRxiv2022.11.07.515477.

Quartarone A, Cacciola A, Milardi D, Ghilardi MF, Calamuneri A, Chillemi G, Anastasi G & Rothwell J (2020). New insights into cortico-basal-cerebellar connectome: clinical and physiological considerations. Brain : a journal of neurology 143, 396–406.

Quoilin C, Fievez F & Duque J (2019). Preparatory inhibition: Impact of choice in reaction time tasks. Neuropsychologia 129, 212–222.

Rossi S et al. (2021). Safety and recommendations for TMS use in healthy subjects and patient populations, with updates on training, ethical and regulatory issues: Expert Guidelines. Clinical Neurophysiology 132, 269–306.

Rossi S, Hallett M, Rossini PM & Pascual-Leone A (2009). Safety, ethical considerations, and application guidelines for the use of transcranial magnetic stimulation in clinical practice and research. Clinical Neurophysiology 120, 2008–2039.

Rossini PM et al. (2015). Non-invasive electrical and magnetic stimulation of the brain, spinal cord, roots and peripheral nerves: Basic principles and procedures for routine clinical and research application. An updated report from an I.F.C.N. Committee. Clinical Neurophysiology 126, 1071–1107.

Rossini PM, Barker AT, Berardelli A, Caramia MD, Caruso G, Cracco RQ, DimitrijeviImage MR, Hallett M, Katayama Y, Lücking CH, Maertens de Noordhout AL, Marsden CD, Murray NMF, Rothwell JC, Swash M & Tomberg C (1994). Non-Invasive Electrical and Magnetic Stimulation of the Brain, Spinal Cord and Roots: Basic Principles and Procedures for Routine Clinical Application. Report of an IFCN Committee. Electroencephalography & Clinical Neurophysiology 91, 2198–2208.

Rosso C, Perlbarg V, Valabregue R, Obadia M, Kemlin-Méchin C, Moulton E, Leder S, Meunier S & Lamy J-C (2017). Anatomical and functional correlates of cortical motor threshold of the dominant hand. Brain Stimulation 10, 952–958.

Rothwell JC, Hallett M, Berardelli A, Eisen A, Rossini P & Paulus W (1999). Magnetic stimulation: motor evoked potentials. The International Federation of Clinical Neurophysiology. Electroencephalography and clinical neurophysiology Supplement 52, 97–103.

Rurak BK, Rodrigues JP, Power BD, Drummond PD & Vallence AM (2021). Test Re-test Reliability of Dual-site TMS Measures of SMA-M1 Connectivity Differs Across Inter-stimulus Intervals in Younger and Older Adults. Neuroscience 472, 11–24.

Spieser L, Aubert S & Bonnard M (2013). Involvement of SMAp in the intention-related long latency stretch reflex modulation: A TMS study. Neuroscience 246, 329–341.

Stokes MG (2005). Simple Metric For Scaling Motor Threshold Based on Scalp-Cortex Distance: Application to Studies Using Transcranial Magnetic Stimulation. Journal of Neurophysiology 94, 4520–4527.

Stokes MG, Barker AT, Dervinis M, Verbruggen F, Maizey L, Adams RC & Chambers CD (2013). Biophysical determinants of transcranial magnetic stimulation: Effects of excitability and depth of targeted area. Journal of Neurophysiology 109, 437–444.

Stokes MG, Chambers CD, Gould IC, English T, McNaught E, McDonald O & Mattingley JB (2007). Distance-adjusted motor threshold for transcranial magnetic stimulation. Clinical Neurophysiology 118, 1617–1625.

Tokuno H & Nambu A (2000). Organization of nonprimary motor cortical inputs on pyramidal and nonpyramidal tract neurons of primary motor cortex: An electrophysiological study in the macaque monkey. Cerebral cortex (New York, NY : 1991) 10, 58–68.

Vassiliadis P, Derosiere G, Grandjean J & Duque J (2020). Motor training strengthens corticospinal suppression during movement preparation. J Neurophysiol 124, 1656– 1666.

Vassiliadis P, Grandjean J, Derosiere G, de Wilde Y, Quemener L & Duque J (2018). Using a double-coil TMS protocol to assess preparatory inhibition bilaterally. Frontiers in Neuroscience 12, 1–14.

Vesia M, Barnett-Cowan M, Elahi B, Jegatheeswaran G, Isayama R, Neva JL, Davare M, Staines WR, Culham JC & Chen R (2017). Human dorsomedial parieto-motor circuit specifies grasp during the planning of goal-directed hand actions. Cortex 92, 175–186.

Wang Y, Cao N, Lin Y, Chen R & Zhang J (2020). Hemispheric Differences in Functional Interactions Between the Dorsal Lateral Prefrontal Cortex and Ipsilateral Motor Cortex. Frontiers in Human Neuroscience 14, 1–6.

Wilhelm E, Quoilin C, Derosiere G, Paço S, Jeanjean A & Duque J (2022). Corticospinal Suppression Underlying Intact Movement Preparation Fades in Parkinson’s Disease. Mov Disord 37, 2396–2406.

Zrenner C, Desideri D, Belardinelli P & Ziemann U (2018). Real-time EEG-defined excitability states determine efficacy of TMS-induced plasticity in human motor cortex. Brain Stimulation 11, 374–389.

